# Deciphering the tissue specificity of the protein secretory pathway in humans

**DOI:** 10.1101/070870

**Authors:** Amir Feizi, Francesco Gatto, Mathias Uhlen, Jens Nielsen

## Abstract

Proteins that are components of the secretory machinery form a cellular pathway of paramount importance for physiological regulation, development and function of human tissues. Consistently, most secretory pathway components are ubiquitously expressed in all tissues. At the same time, recent studies identified that the largest fraction of tissue-specific proteins consists of secreted and membrane proteins and not intracellular proteins. This suggests that the secretory pathway is distinctively regulated in a tissue-specific fashion. However, a systematic analysis on how the protein secretory pathway is tuned in different tissues is lacking, and it is even largely unexplored if the secretome and membrane proteome differs in, for example, posttranslation modifications across tissues. Here, analyzing publically available transcriptome data across 30 human tissues, we discovered the expression level of key components previously categorized as housekeeping proteins were specifically over-expressed in a certain tissue compared with the average expression of their corresponding secretory pathway subsystem (e.g. protein folding). These *extreme genes* define an exceptional fine-tuning in specific subnetworks, which neatly differentiated for example the pancreas and liver from 30 other tissues. Moreover, the subnetwork expression tuning correlated with the nature and number of post translational modification sites in the pancreas or liver-specific secretome and membrane proteome. These patterns were recurrently observed also in other tissues, like the blood, the brain and the skeletal muscle. These findings conciliate both the housekeeping and tissue-specific nature of the protein secretory pathway, which we attribute to a fine-tuned regulation of defined subnetworks in order to support the diversity of secreted proteins and their modifications.

## Introduction

The protein secretory pathway in eukarya involves an elaborate integration of functional modules compartmentalized in the endoplasmic reticulum (ER) and Golgi apparatus. These modules are responsible for stepwise post-translational modifications (PTMs) (such as folding and glycosylation) and transport of secretory proteins(Fig. 1A). The essence of this intricate cell machinery is to guarantee the functionality of the proteins routed to the extracellular space, either as membrane or secreted proteins. A functioning secretory pathway is essential for human body physiology. The majority of hormones and enzymes produced by the endocrine and exocrine systems, but also receptors and channels, anchors and extracellular matrix components, coagulation factors, and other molecular transporters are all clients of the secretory pathway. Given these housekeeping roles, the pathway is considered to be expressed in most human tissues. Unsurprisingly, dysfunction of the secretory pathway is causally or indirectly implicated in a variety of systemic or developmental diseases, like cancer, diabetes, Parkinson’s disease, and congenital neurodegenerative disorders (Freeman, 2001; Pohlschröder et al, 2005; Sherwood, 2015; Uhlén et al, 2015). Over the past 40 years, we have acquired an exhaustive picture of the molecular components (i.e. proteins) of the secretory pathway and its clients, i.e. the secretome and the membrane proteome (Lippincott-Schwartz, 2011; Novick et al, 1981; Novick et al, 1980; Rothman, 2014; Südhof & Rothman, 2009). However, until recently, a systemic genome-scale map of the protein secretory pathway in eukarya was not available (Feizi et al, 2013). Therefore, the study of regulation of the protein secretory pathway at the system level is largely unexplored.

**Figure 1.**
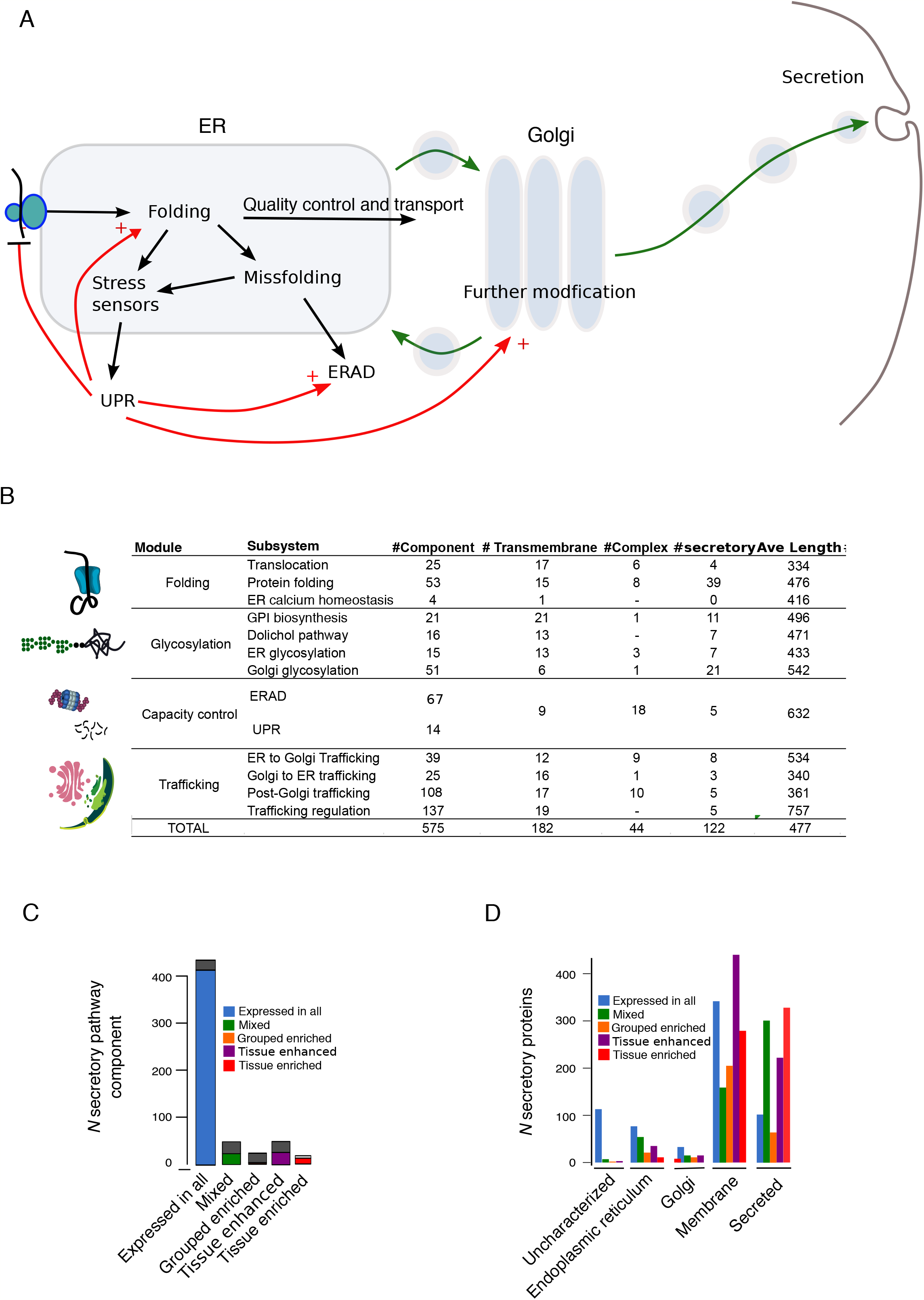
Expression of the secretory pathway and its clients across humantissues. **(A)** the schematic representation of the core secretory pathway functional modules. The black arrow indicates processes related with processing proteins while arrows with red and green shows in order the unfolded protein response(UPR) targets and transport steps (B) The properties and functional modules of the reconstructed human secretory pathway, for a total of 575 components. (C) The expression categories of human secretory pathway genes based on GTEx and HPA data. (D) The expression categories of human secreted or membrane protein-encoding genes based on GTEx and HPA. Only protein with N-terminal signal peptide are included. Proteins were grouped according to localization.

In human tissues, tissue-specific features are generally the result of complex regulatory cascades through development stages, which are aimed to specifically tune the expression of the different cellular processes. In the case of secretory and membrane proteins, which are the clients of the secretory pathway, the differences across tissues are fairly obvious (e.g. insulin for pancreas as opposed to renin for kidney). Considering the above-mentioned housekeeping character of the secretory pathway itself, these tissue-specific differences should arise from an elaborate fine-tuning in the expression of the pathway components. However, it is still unclear how different tissues regulate the recently mapped secretory pathway, nor if this regulation correlates with the different needs of tissue-specific membrane and secreted proteins, in terms of processing, transport, or specific PTMs. Compensating for this knowledge gap could result in three valuable outcomes: first, to provide a rational approach to study the diseases linked with secretory pathway by a detailed mechanistic mapping of all pathway steps in different tissues; second, to aid engineering of biopharmaceutical protein production, since the current bottleneck in the production of human proteins is the functional difference between the host (e.g. CHO cells) and parent secretion system (Golabgir et al, 2016); Kildegaard et al, 2013); and finally to advance our understanding on the role of gene expression in the development of tissue-specific function in humans.

Here we performed a systematic analysis of gene expression for the 575 core components of the secretory pathway across 30 human tissues, focusing on tissue-specific fine-tuning and its relation with the requirements for each tissue-specific secretome and membrane proteome.

## Results and Discussion

### Is the secretory pathway a housekeeping cell machinery?

The secretory pathway is an essential and ubiquitous machinery in human cells, and yet provides specific functionality depending on the tissue. This is evident in secretory tissues belonging to the endocrine system (e.g. the pancreas), but it applies more generally to most human tissues. To dissect the extent to which the expression level of secretory pathway components is tuned in the different tissues as opposed to ubiquitous expression in a housekeeping fashion, we investigated the tissue-wise variation in the mRNA levels of the corresponding genes. We used two independent and comprehensive RNA-seq datasets of 30 human tissues, the Genotype-Tissue Expression Project (GTEx, (Melé et al, 2015)) as the main set and the Human Protein Atlas (HPA, (Uhlén et al, 2015)) as a validation set. Both datasets achieved unprecedented resolution on tissues’ RNA levels with significant between-study correlation (Uhlén et al, 2016). We opted to use the GTEx dataset as the main set because it included brain sub-regions and blood and featured a higher number of replicates and sequencing depth. These studies adopted expression categories to classify tissue-specificity of genes (e.g. “tissue elevated”, if the gene mRNA levels were at least five-fold higher in a particular tissue as compared to all other tissues).

Next, starting from the yeast secretory model (Feizi et al, 2013), we reconstructed a generic human secretory pathway network of 575 core components (see Materials and methods), which were allocated to 13 specific functional modules (Fig. 1B). Of all 575 secretory pathway components, ~83% (n=478) of the genes belonged to the same expression category according to both GTEx and HPA (Fig. 1B). Of these 478 genes, ~86% (n=414) belonged to the “expressed in all” category and ~13% (n=64) were in tissue-specific categories (such as “tissue enriched” and “group enriched”) (Fig. 1C-EV1). The 414 “expressed in all” genes showed similar expression distribution (in terms of log_10_ FPKM) across tissues both in GTEx and HPA (median expression ~=10 FPKM), yet tissues such as pancreas, skeletal muscle, heart and liver displayed slightly lower median expression compared to other tissues (Fig. S1). Collectively, this is consistent with the notion that the secretory pathway is a housekeeping machinery.

On the other hand, it has been previously shown that in most tissues 10-20% of the transcriptome translates into secreted or cell-membrane proteins (Uhlén et al, 2015). This fraction increases up to 70% in secretory tissues such as pancreas and salivary-glands. Most strikingly, the largest fraction of tissue-specific proteins are secreted or membrane proteins as opposed to intracellular proteins (Fig. 1D) (Uhlén et al, 2015). These facts argue that the secretome and the membrane proteome are the single most defining class of tissue-specific proteins in any human tissue. Based on HPA data, 5,670 proteins were predicted to be uniquely secreted or membrane proteins, among which 3,328 were predicted with a N-terminal signal peptide that dictates the entering and passing through the secretory pathway for harboring proteins. Of these 3,328 proteins, 1,218 were secreted proteins and 1,607 were plasma membrane proteins. Most of these were assigned to the tissue-specific categories (e.g. “tissue enriched”) (Fig. 1D). The remaining 503 proteins were localized in the lumen or the membranes of the ER, Golgi or other organelles (Fig. 1D). Among the 1,242 proteins without predicted signal peptide, there were 680 proteins predicted to be secreted by unconventional secretion (secretome P NN-score > 0.6) among which 12 belonged to the secretome and 507 to the membrane proteome (Bendtsen et al, 2004; Nickel& Seedorf, 2008) (EV2). Considering this high degree of tissue-specificity of secreted proteins, we contended that the secretory pathway should be regulated differently across tissues, despite its ubiquitous expression. Indeed, the brain and the pancreas, for example, differ in secreted and membrane proteins both in terms of protein function and physiology, and we deemed this unlikely to be solely represented by the small fraction (13%) of “tissue elevated” genes in the pathway.

To assess within-tissue variation, we correlated the expression profile of proteins belonging to the secretory pathway, but limited to the 414 “expressed in all” genes across 30 tissues (Fig. 2). Hierarchical clustering of the correlation coefficients supported on one hand a housekeeping role of the pathway, in that most tissues showed medium to high correlation with each other (ρ = 0.83 to 0.98), despite the presence of sub-clusters with higher in-between correlation (e.g. between uterus, cervix, vagina). On the other hand, nine tissues from the liver, pancreas, blood, kidney, skeletal muscle, heart, testis, and brain (cerebellum, and cerebrum) formed isolated clusters, with low to medium correlation with the tissues in the larger cluster (median coefficient *ρ* = 0.57 +-0.17, permutation test *p* < 0.05). This clustering pattern partially was observed also using the HPA data, with the difference that the thyroid clustered instead with the salivary gland (missing in GTEx) and pancreas clustered with the skeletal muscle, albeit presenting an even smaller correlation with other tissues compared to GTEx (Fig. S2). This is suggestive that even ubiquitously expressed secretory pathway genes display a degree of tissue-specific modulation.

We observed that at the transcriptome level, the pancreas, pituitary gland, and blood in GTEx and the pancreas, bone marrow and salivary gland in HPA clustered away, with negligible effect of including the secretory pathway genes, warranting a potential confounding effect due to a generally deviance of their expression profile (Fig S3). Collectively, these results indicate the secretory pathway mostly featured “expressed in all” genes but these can be further modulated in a tissue specific-function. We therefore evaluated whether this fine-tuning in housekeeping genes reflected a specific functional process important for a certain tissue by analyzing secretory pathway subsystems.

### Expression levels of the secretory pathway subsystems are tuned in each tissue

In our previous study for yeast (Feizi et al, 2013), we defined 169 secretory pathway components that were mapped in to functional modules (subsystems) representing distinct functional modules such as translocation, protein folding, ER glycosylation etc. Using the same approach (see Methods), we allocated the 575 core components of the human secretory pathway into these same 13 subsystems (Fig. 1A) and (B) , Table EV1). We sought to investigate whether in the outlier tissues the 414 “expressed in all” genes were fine-tuned uniformly across the spectrum of functions in the secretory pathway or preferentially in a specific subsystem(s), hinting that these tissues need to adjust housekeeping genes according to tissue-specific requirements. We started by analyzing first the pancreas, one of the least correlated tissues (median ρ = 0.49 +-0.11). We calculated for each subsystem the correlation between the expression levels in the pancreas versus any other tissue (Fig. 3A). Most subsystems correlated fairly well (ρ > 0.6, p < 0.05), while others fluctuated substantially to very low to high correlations according to the compared tissue (e.g. translocation). The correlation coefficients displayed a broad dynamic range between the lowest (translocation, ρ = 0.03) and highest value (GPI-biosynthesis, ρ = 0.94). HPA data yielded highly consistent results (Fig. S5). This is suggestive that even if pancreas was a relative outlier when considering the bulk of housekeeping components of the secretory pathway, at the subsystem level, the pancreas had a relatively similar profile to many other tissues for many secretory functions, but it severely differed in terms of expression of proteins involved in translocation and trafficking subsystems. For example, the expression of GPI biosynthesis module correlated between the brain cerebrum and the pancreas, but no correlation was observed for translocation (Fig. 3B). Consistently, the pancreas and the liver showed strong correlations in most subsystems, therefore they are expected to share similar functionality of the secretory pathway in general, as suggested by our results above (Fig. 2). The example of pancreas clearly highlights that “expressed in all” genes undergo tissue-specific adjustment geared towards specific functional modules rather than fine-tuning the pathway as a whole.

**Figure 2.**
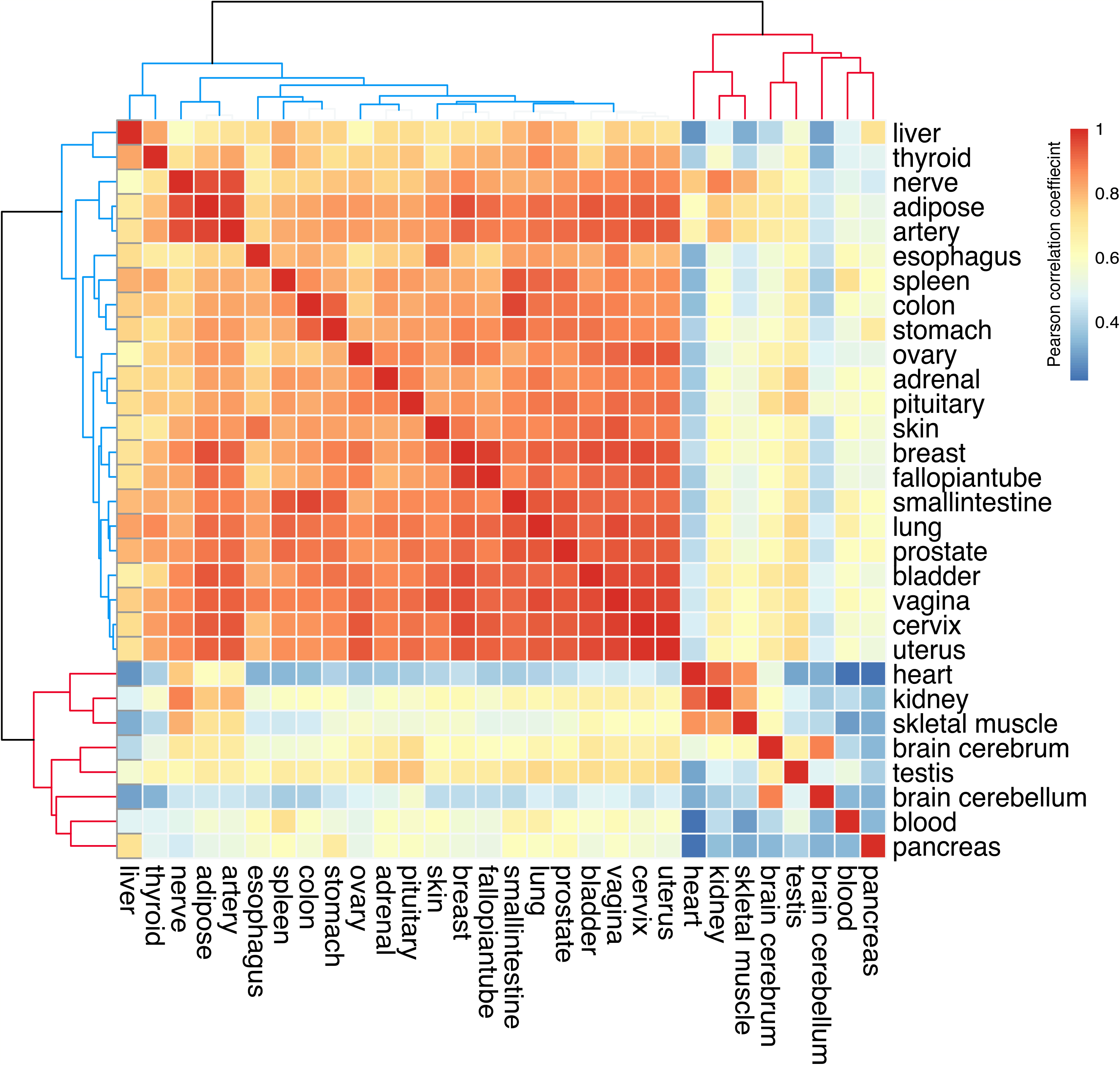
Hierarchical clustering of between-tissue correlation coefficients for the expression of secretory pathway genes in the “expressed in all” category. Heatmap of the hierarchical clustering of 30 human tissue pairwise correlation (Pearson correlations) using the expression profiles of their secretory pathway genes belonging to “expressed in all” category. The subclusters discussed in the text body are shown with distinct colors for the dendrogram clades.

**Figure 3.**
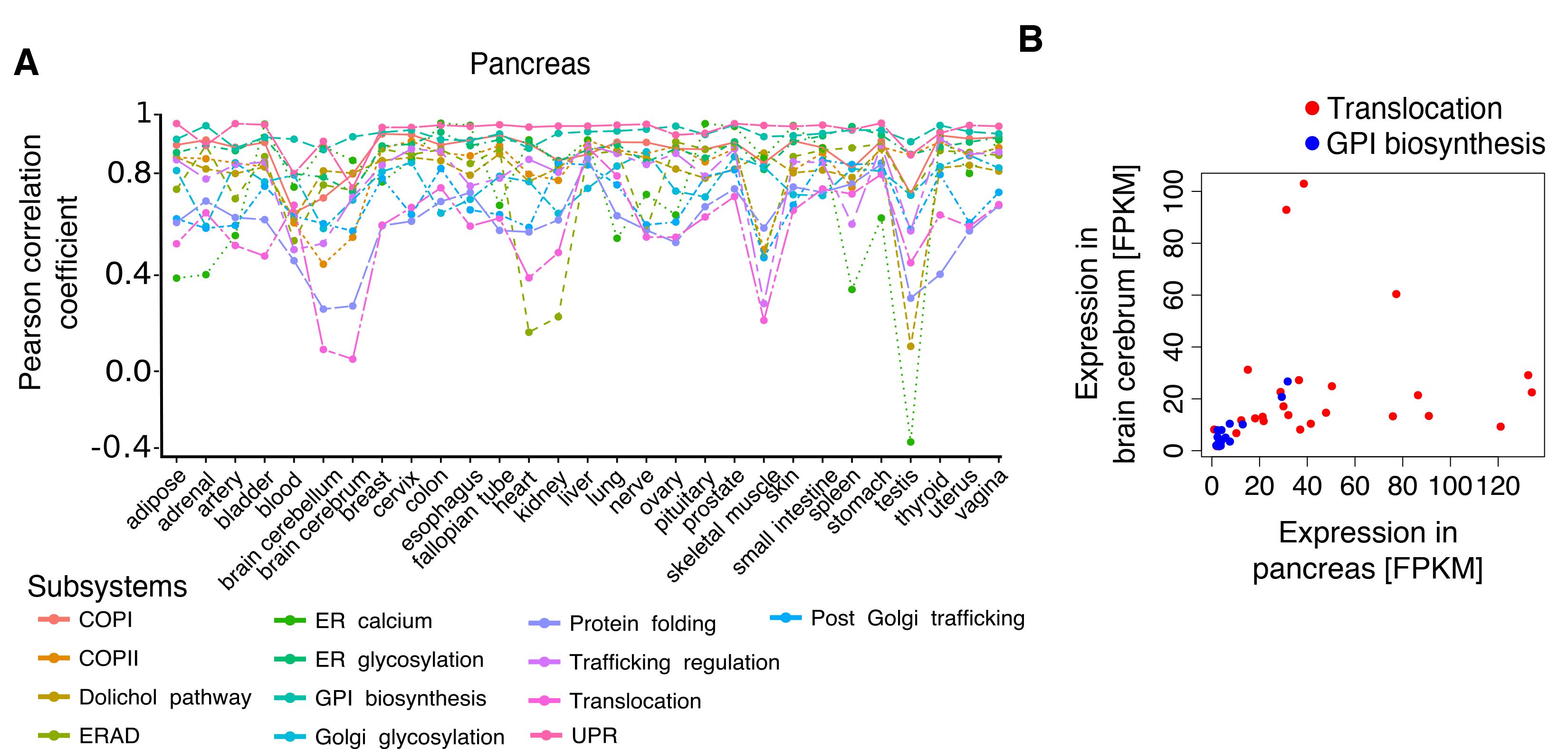
Correlation of the expression in pancreas vs. other tissues for “expressed in all” genes grouped by secretory pathway subsystems. (A) Multi-line plot for Pearson correlation coefficients of the gene expression profiles of each subsystem in pancreas versus other tissues. Only “expressed in all” genes were included. Each line represents a different subsystem. (B) Correlation between the expression of “expressed in all” genes in the translocation (red dots) or GPI-biosynthesis (blue dots) subsystems between the cerebrum and the pancreas.

We next generalized the results from pancreas to the other tissues by performing a correlation analysis of all tissue-pairs (n=11700) for each subsystem, limited to “expressed in all” genes. Interestingly, subsystems could be seen as either extremely correlated across most tissue-pairs, for example UPR (unfolded protein response), or with wider correlation coefficient distributions, like trafficking regulation (Fig. 4A).This indicates that certain subsystems might have tissue-specific gene expression modulation. Therefore, we sought to explore the tissue-pairs with non-significant subsystem correlation (ρ < 0.6). To identify the most tissue-specific gene among the “expressed in all” genes in each subsystem, we run Grubbs test (Grubbs, 1950) using both GTEx and HPA data and we picked the overlapped detected outliers, assuming that the total expression level of each subsystem vary among tissues (see Materials and Methods). As result, we discovered that within each subsystem tissues, there were a set of extreme genes (i.e. outliers) that were specific or shared between few tissues despite defined as expressed in all (Fig. 4B), These genes were likely responsible for the low tissue-pair correlations in some subsystems uncovered above (Fig. 4A, given that none of these extreme genes belonged for example to UPR, which was the most correlated subsystem across tissues (Fig. 4B). Noteworthy, these extreme genes still belong to the “expressed in all” category, meaning that they are expressed in all tissues, however in a specific (or few) tissue they displayed exceptionally high expression compared to the average expression of the subsystem they were part of. Most extreme genes belonged to post-Golgi trafficking or trafficking regulation subsystem (Fig. 4E). This is consistent with the previous result that low tissue-pairs correlations were observed particularly in these subsystems for “expressed in all” genes (Fig. 4A). On the other hand, brain cerebrum has the highest number of extreme genes (n=17) that five of them are unique or shared up to seven tissues (Fig. 4B and 4F). For example, focusing on extreme genes in the skeletal muscle and the pancreas, as two tissues with lowest correlations (Fig. 4B), extreme genes that were uniquely associated with either of these tissues showed an evident higher expression level comparing to other tissues (for example OPTN for skeletal muscle or SEL1L for pancreas). Although, some of the extreme genes were shared between tissues like RAB10 (skeletal muscle and esophagus) and PDIA4 (pancreas, liver and kidney), the expression level was always higher for one of the tissues (Fig. 4D) and (4E).

**Figure 4.**
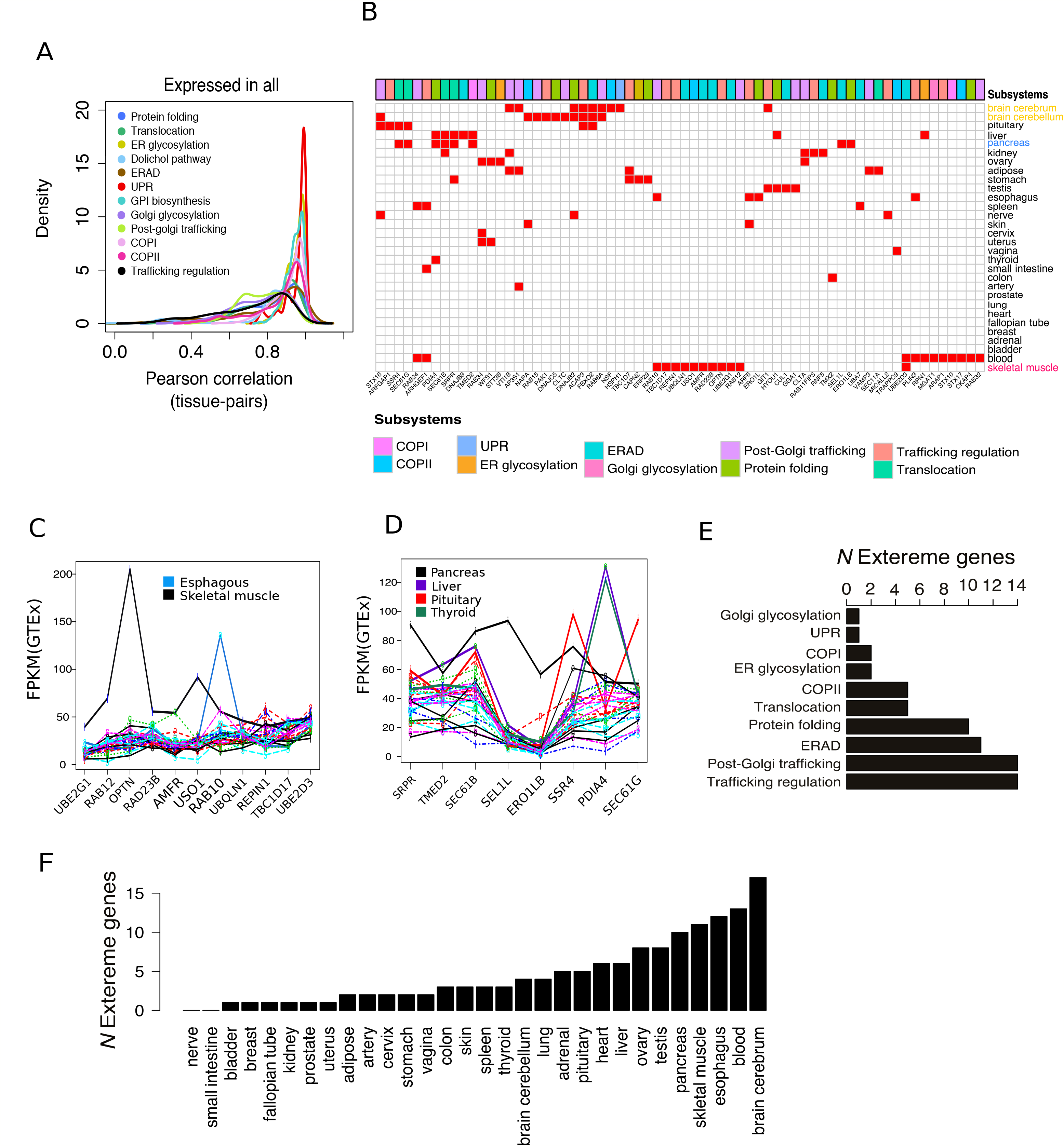
Detecting the tissue specific activity of secretory pathway genes “expressed in all” tissues. (A) The kernel density plot of tissue-pairs correlation coefficients calculated for genes in each subsystem assigned to the “expressed in all” category. (B) The binary heat map for the detected tissue-specific extreme genes (columns) in each subsystems of the secretory pathway across 30 tissues (rows). Some tissues and their corresponding extreme genes are shown with the same color. The total number of detected extreme genes per subsystem is summarized as horizontal bar plot next to the heatmap. (C,D) Multi-line plot of the FPKM values for the extreme genes in the skeletal muscle and the pancreas (black lines). Each line represents a tissue. (E,F) The number of the detected extreme genes per subsystems and each tissue.

Since the analysis above only contained “expressed in all” genes, we controlled that no biases in the above correlations could be imputed to low subsystem size or exclusion of genes in tissue-specific categories in the different subsystems. As suggested by the fact that 86% of pathway genes assigned to a consensus category were “expressed in all”, the greatest fraction of all subsystems also belonged to the “expressed in all” genes (Fig S6A). For instance, translocation genes were all assigned to “expressed in all” category, while certain subsystems, like ERAD (endoplasmic reticulum-associated degradation) and protein folding featured a small fraction of genes in tissue-specific categories. These sum up to 12 genes, which were all testis-specific proteins except CRYAA, which was a kidney-specific chaperone (Fig. S6B). These genes in testis belonged to ERAD (5 genes), protein folding (2), Golgi glycosylation (2) and trafficking regulation (2). Therefore, we ruled out that neglecting these genes in tissue-specific categories contributed to explain the lower correlation in the expression of the secretory subsystems observed for some tissues. This strengthens the hypothesis that subsystems experience expression fine-tuning in a tissue-specific fashion predominantly in the case of “expressed in all” genes. Finally, we tested the between-tissue correlations limited to non-“expressed in all” genes (n =64, based on GTEx) at the subsystem level. The distribution of tissue-pair correlations spanned a broad range of coefficient values in the subsystems with genes in tissue-specific categories (such as trafficking subsystems, Fig. S7). Thus, some tissue-pairs were lowly correlated in a given subsystem, suggesting that these tissue-specific genes contribute (albeit modestly) rendering these subsystems tissue-specific. The same conclusion cannot hold for subsystems such as translocation or ER glycosylation, which featured exclusively “expressed in all” genes.

These findings show that even if secretory pathway genes are expressed rather consistently in all tissues, individual tissues can spike the expression of selected genes in defined subsystems in a tissue-specific fashion. This indicates that the secretory pathway is a housekeeping machinery, but certain functional modules, particularly within protein folding and trafficking, need to be finely adjusted according to tissue-specific requirements. Next, we explored if this tissue-specific expression tuning correlated with requirements for PTMs and functions of membrane proteins or proteins secreted that are typically specific to the different human tissues.

### Tissue specific properties of human secretome and membrane proteome dictates the expression tuning of the secretory pathway components

So far there has not been a systemic approach on how the secretory pathway is regulated in the different tissues to provide the known degree of specificity in their secretome and membrane proteome. This has also been hindered by the fact that it has not yet been elucidated if and how the properties of the secretome and membrane proeome differ across human tissues. We hypothesized that the tissue-specific fine-tuning of certain secretory pathway subsystems is associated with the properties of the proteins that are client of the pathway in a certain tissue. To investigate this, we analyzed the tissue-specific properties of the secretome and membrane proteome.

As already discussed earlier in (Fig. 1C), we assembled a list of 4,098 of conventional (with signal peptide) (n=3,328) and unconventional (without signal peptide) (n=680) secreted or membrane proteins. We obtained the GTEx expression profiles of the associated genes and performed hierarchical clustering of tissues based on their expression correlation matrix (Pearson correlation). We focused on genes in tissue-specific categories (2,047 genes) and used the scaled expression instead of FPKM for gaining better resolution. We observed a neat separation of “tissue-enriched” genes for the corresponding tissue (Fig. 5). The liver (N secreted or membrane proteins = 90) and the pancreas (N=30) showed the highest ratio of secreted to membrane "tissue-enriched” genes. The testis (101), blood (34), kidney (25), and brain-cerebrum (21) displayed the highest ratio of membrane to secreted “tissue-enriched” genes (“localization” in (Fig. 5).

**Figure 5.**
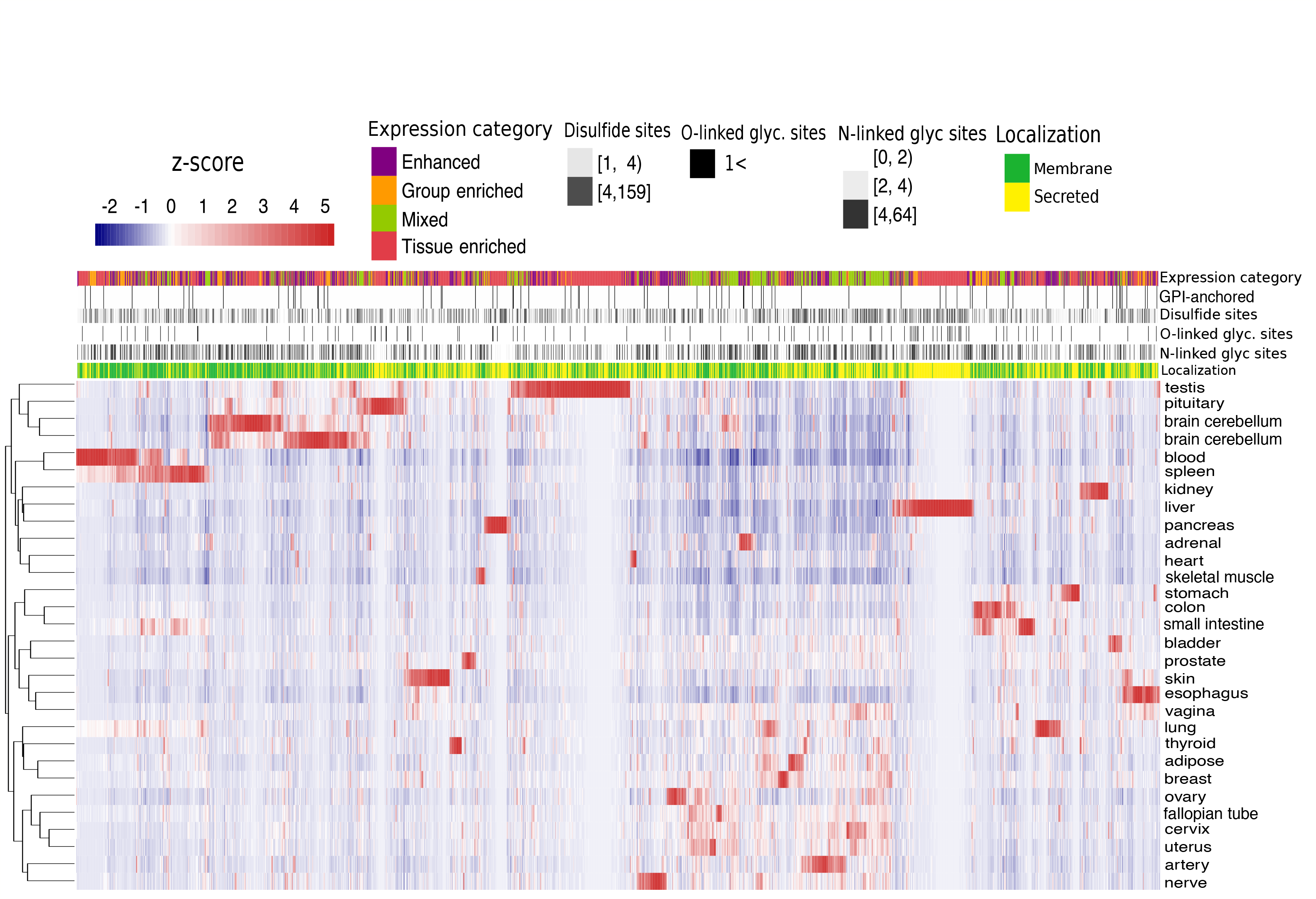
Hierarchical clustering of human tissue specific secretome and membrome expression and their properties. Each row represents a tissue and each column a tissue-specific secreted or membrane protein-encoding gene. Z-scores are scaled FPKM values for the expression of a given gene in a given tissue versus other tissues. The annotation bars above the heatmap provide information regarding the expression category, number of disulfide, N-linked or O-linked glycosylation sites, GPI-anchored sites and localization. The number of the sites are discretized to separate low, medium and high number of PTM sites.

Secretory proteins, unlike intracellular proteins, undergo specific modifications steps through the ER and Golgi that guarantee their function once exposed to the extracellular environment. Of the secretory PTMs, glycosylation, sulfation and glycosylphosphatidylinositol (GPI anchor) are the main modifications. The secretome and membrane proteome are highly tissue-specific, and therefore should impose specific PTMs processing within the secretory pathway depending on the tissue. This might correlate with the fine-tuning of specific secretory subsystems attributed to extreme genes that we observed before. To investigate PTMs differences, we obtained information from UniProt on number of sites for N-glycosylation (NG), disulfide (DS), O-glycosylation (OG), GPI-anchored (GP) for secreted and membrane proteins specific to each tissue, and integrated this information with the clustering result (Fig. 5). In general, most of the tissue specific secreted and membrane proteons are enriched with *N*-linked glycosylation and disulfide sites, however, pancreas and pituitary are two tissues which stand out as they have less enrichment in the *N*-linked glycosylation sites in their secretome while they are highly enriched in disulfide sites (Fig. 5). O-linked and GPI-anchored sites are rather tissue-specific. For example, liver secreted proteins are enriched in O-linked, whereas brain sub-regions are enriched with GPI-anchored membrane proteins. In the following we interpreted these results in respect of the similarity between the tissues secretory function and if there are any correlation with identified extreme genes.

### Pancreas, liver and kidney

Both in the liver and the pancreas, genes in tissue-specific categories were mainly secreted protein (Fig. 5). In the pancreas, highly expressed genes were CELA2A, CTRB2 (peptidase), CTRB1, insulin (INS), PNLIP (pancreas lipase), and AMY2A (alpha amylase); in the liver, some of these genes were F9, C8A, or CFHR2, which encode proteins involved in plasma complement binding or lipid transporters (see EV3 for the full list). These proteins harbored multiple glycosylation and disulfide sites, which require plenty of glycan, energy and folding processing units within the secretory pathway of these tissues. Also, as most of the proteins were secreted, with continues outflux to the extracellular space, therefore we expected a high pressure on transport subsystems. Collectively, these properties should require specific modulation of the folding, glycosylation and transport system in these two tissues. This argument was in line with the function of the extreme genes previously detected for pancreas and liver (Fig. 4B). Strikingly, 4 of these were overlapping in these two tissues, *PDIA4 TMED2, SRPR, SEC61B* ((Fig. 4B). which belonged to the folding and transport subsystems. Interestingly, it was earlyer reported that the expression level of extreme genes *ERO1LB*, and *SEL1L*, foldases with strong disulfide isomerase activity, was pancreas-specific (Tufo et al, 2014). This correlates with the putative exceptionally high flux of secreted proteins with disulfide sites (Fig. 5). On the other hand, we considered *PDIA4* as an interesting example of tissue-specific fine-tuning of a secretory pathway subsystem, because it was shared as an extreme gene by the pancreas, the liver and the kidney (with higher expression in kidney and liver). These tissues featured among the highest numbers of over-expressed tissue-specific secreted or membrane proteins enriched with disulfide bonds (Fig. 6B). Consistent with processing of disulfide bonds, *PDIA4* was expressed in all tissues, but the expression level of this isoenzyme was significantly higher in these three tissues (Fig. 6B). As a second example, *SRPR, SSR4, TMED2, SEC61B*, and *SEC61G* are involved in the translocation and trafficking subsystems. *TMED2* is an established specific protein for the pancreas (Blum et al, 1996), and has been shown to play a critical role in cargo detection from ER (COPII vesicle-mediated) and to regulate exocytic trafficking from the Golgi to the plasma membrane (Dominguez et al, 1998; Luo et al, 2011; Stepanchick & Breitwieser, 2010).

**Figure 6.**
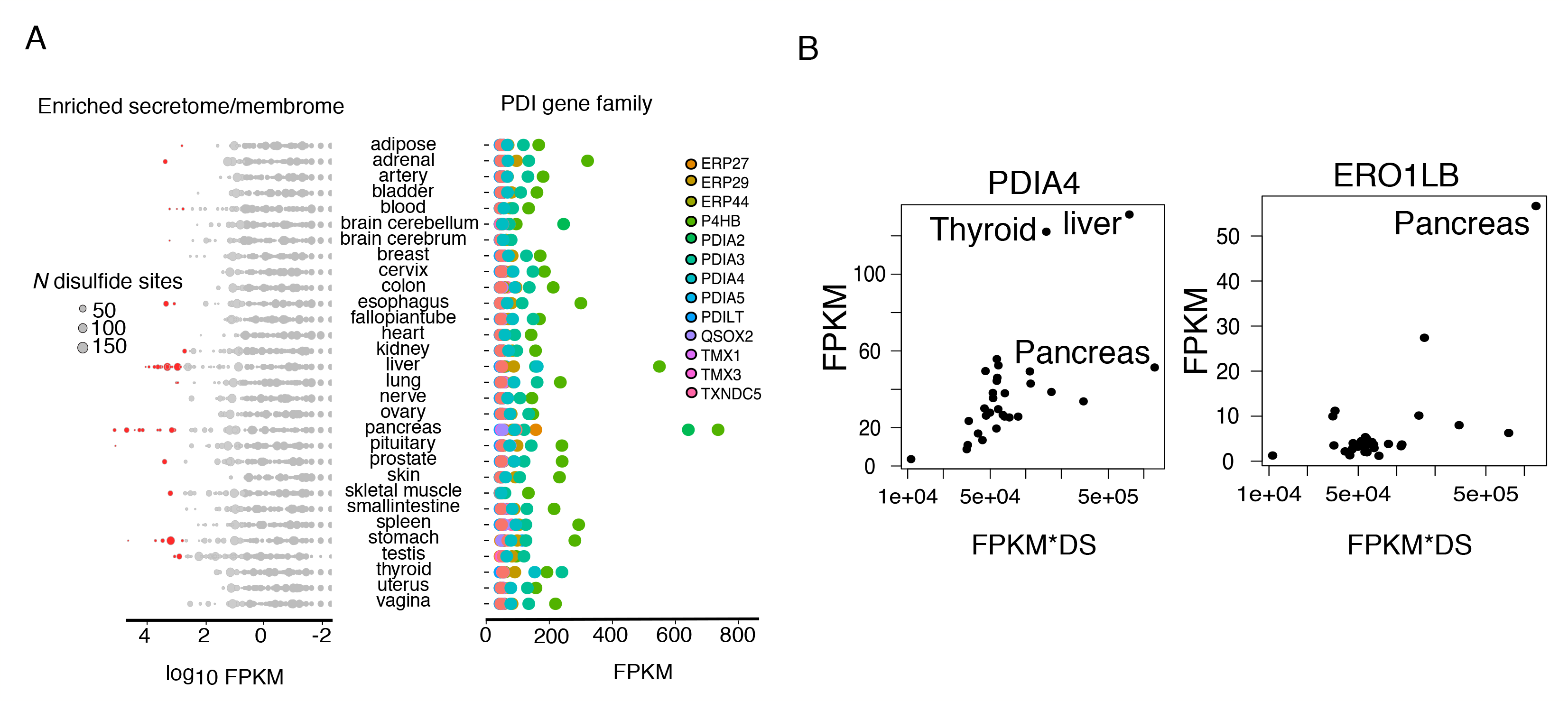
Tissue-wise expression of the tissue-specific secretome and membrome and correlation between their number of disulfide bonds and expression of secretory pathway components in the PDI gene family. **(A)** The double side horizontal dot plots summarize the expression level of the tissue-specific secretome and membrome in each tissue(left side) and the expression level of the PDI gene family(right side), foldases and isomerases responsible for the proper folding of proteins with disulfide bonds. The number of disulfide bonds for each protein is proportional to the dot size. Expression is in log_10_ FPKM and genes with FPKM > 1000 were shown in red. (B) The scatter plots depicting the expression values of the *PDIA4* and *ERO1LB* in 30 tissues against the total sum of the FPKM values of the tissue-enriched secreted proteins multiplied by the corresponding number of the disulfide bonds of each encoded proteins.

Therefore, tuning the expression level of specific gene(s) contribute to the tissue specialization of the secretory pathway function in connection with enriched specific PTMs load or by running the transport system efficiently. We therefor suggest tissue-specific roles for most of the detected extreme genes in liver, pancreas and kidney.

### Brain - the cerebrum, the cerebellum and the pituitary gland

Focusing on the secretome of brain sub-regions, the cerebrum, the cerebellum and the pituitary gland clustered together and yet displayed an evident degree of specificity for their corresponding secreted and membrane proteins (Fig. 5). Most brain enriched proteins, especially for the cerebrum and the cerebellum, were membrane proteins (Fig. 5). Most of brain tissues enriched proteins had multiple N-glycosylation and/or disulfide sites (Fig. 5). Correspondingly, among the extreme genes detected for brain sub-regions, for instance, *FBXO2* and *ACAP3* were shared extreme genes across the three sub-regions. *ACAP3* is presumed to be a GTPase activator and *FBXO2* is presumed to recognize and bind denatured glycoproteins (in the ERAD pathway), preferentially those of high-mannose type, which are strongly represented among brain membrane proteins. Considering that dysregulation in proper O-mannosylation in brain has been linked with serious congenital diseases (Haltiwanger& Lowe, 2004), we speculate that *FBXO2* could play a key role in the quality control of brain membrane protein mannosylation. The cerebellum had four exclusive extreme genes, belonging to post-Golgi trafficking *(CLTC, RAB15)*, trafficking regulation *(PAK1)* and protein folding *(DNAJC5)*. Interestingly, in the pituitary gland, an active endocrine tissue, we detected two extreme genes shared with the pancreas which both belong to the translocation subsystem *(SSR4* and *SEC61G)*. As both tissues have high flux of secretory proteins and high expression of these proteins may play an important role in enabling a high translocation rate of hormones from the cytoplasm to the ER.

### Testis

The testis had the highest number of tissue-specific proteins that are client of the secretory pathway (n=172), mostly located on the membrane (n=113) (Fig. 5). These proteins featured multiple disulfide and/or N-glycosylation sites. Most of these proteins (58%) enriched GO terms are associated with testis-specific processes, like spermatogenesis and sperm-egg recognition (Fisher’s exact test *p* <0.001). Genes including *GGA1, CUL1, HYOU1* and *GIT1* were detected as extreme genes in testis. *GGA1* encodes a member of the Golgi-localized gamma adaptin ear-containing protein, ARF-binding (GGA) protein family, and regulates the trafficking of proteins between the trans-Golgi network and the lysosome (Boman et al, 2000; Puertollano et al, 2001). The *CUL1* is a component of multiple SCF (SKP1-CUL1-F-box) E3 ubiquitin-protein ligase complexes that mediates the ubiquitination of proteins in cell cycle progression, signal transduction and transcription (Chew et al, 2007; Goldenberg et al, 2004). *HYOU1* encodes a heat shock protein 70 family member. This gene has alternative transcription start sites whose cis-acting segment in the 5' UTR is involved in stress-dependent induction, resulting in the accumulation of this protein in the endoplasmic reticulum (ER) under hypoxia. This accumulation has been suggested to play a pivotal role in protein folding (Meunier et al, 2002; Ozawa et al, 1999). *GIT1* was a shared extreme gene between testis and cerebrum. It has been suggested to serve as a scaffold to convey signaling information which control vesicle trafficking, adhesion and cytoskeletal organization (Manabe et al, 2002). Altogether, these genes seemed to be critical for folding and trafficking subsystems, which is consistent with the notion that the testis featured the highest number of clients of the secretory pathway.

### Blood cells

Blood cells were characterized by 34 membrane and 14 secreted tissue-specific proteins. Strikingly, while none of the membrane proteins have O-linked glycosylation sites, 12 out of the 14 secreted proteins in blood cells carry O-glycosylation sites (Fig. 5). In any other tissue except the liver, very few tissue-specific secreted or membrane proteins had O-glycosylation sites (Fig. 5). Gene Ontology enrichment analysis of these genes identified GO terms related to the defense response to Gram-positive bacteria (Fisher’s exact test *p* < 0.001). Blood cells had 6 exclusive extreme genes that belonged each to a specific subsystem ranging from ER glycosylation, protein folding to post-Golgi trafficking (Fig. 4B). Also, blood cells shared the least number of extreme genes with other tissues, in a analogous fashion as the skeletal muscle. For instance, *MGAT1* is a glycosyltransferase involved in ER glycosylation. *STX10* and *STX17* are soluble N-ethylmaleimide-sensitive factor-attachment protein receptors (SNAREs) involved in vesicular transport from the late endosomes to the trans-Golgi network (Tang et al, 1998). *STX17* has been recently shown to be involved in autophagy through the direct control of autophagosome membrane fusion with the lysosome membrane (Diao et al, 2015), which is consistent with the specialization of defense response of blood cells against bacterial infections.

### Skeletal muscle

The skeletal muscle has a relatively small specific secretome (Fig. 5). Only 12 genes were assigned as secretory proteins, of which 9 were membrane proteins, carrying multiple *N*-glycosylation and few disulfide sites (Fig. 5). The molecular function of these genes is linked to the regulation of ion transport. The low number of secreted or membrane proteins in skeletal muscle seemed at odds with the earlier finding that it as one of the tissues with the highest number of extreme genes (N = 11, (Fig. 4B). Upon closer inspection, the extreme genes with highest expression *(RAB12, OPTN, RAB10*, and *USO1* in Fig 4C) are all involved in vesicle trafficking steps. Among these genes *OPTN* had the highest expression. It has been shown to play an important role in the maintenance of the Golgi complex, in membrane trafficking, in exocytosis, through its interaction with myosin VI and *Rab8* (Sahlender et al, 2005; Vaibhava et al, 2012). To date, no tissue-specific activity has been reported in skeletal muscles for *OPTN*. Interestingly, *RAB12*, another extreme gene in the skeletal muscle, was found to interact with *OPTN* according to a reconstruction of the human protein-protein interaction network (K Sirohi et al, 2013). This concordance provides yet another example on how fine-tuning the expression of a defined subsystem appears to correlate with the properties required by the tissue-specific secretome or membrane proteome.

## Conclusion

Although Uhlen *et al* (2015) has recently showed that secreted and membrane proteins to a large extent are tissue-specific, the secretory pathway itself, as an elaborated and complex machinery responsible for processing and transporting the secretome and membrane proteome, appeared to be ubiquitously expressed, with few components (13%) found to be selectively expressed in certain tissues. This contradiction was addressed in this study, because there are obvious different physiological and morphological pressures in the various human tissues, which demand for tissue-specific functions of the secretory pathway. Gene expression is a dominant form of biological regulation that contributes to confer tissue-specific functionality to diverse cell processes. Signaling pathways, regulatory loops and biological interactions are also important players in defining the tissue specificity of the secretory pathway, yet modulation of gene expression is a fundamental form of regulation which has not been systematically explored before (Keller& Simons, 1997; Rodriguez-Boulan& Nelson, 1989). Our knowledge on regulation of secretion is comprehensive in certain tissue, for example the secretory pathway has been thoroughly studied in the pancreas or the kidney, as tissues specialized for the production and regulation of insulin and renin respectively, two critical hormones in human physiology (Davis& Freeman, 1976; Fu et al, 2013; Itoh et al, 2003; Poy et al, 2004). However, there has not been any study that compare the expression level of the secretory pathway in these tissues as opposed to other tissues, how this might correlate to the properties of tissue-specific secreted and membrane proteins, nor how this extends from individual component to subsystems to the pathway level.

Here, we discovered that while most of the secretory pathway components (86%) were classified to be expressed in all tissues, tissue-specific modulation of these components in the form of extreme gene correlated with the properties of proteins secreted or exported to the membrane specific for each tissues. The idea of using subsystems rather than a gene centric comparison allowed us to find extreme genes for each tissue. With this in mind, a subsystem like translocation which contained only “expressed in all” genes in should not be regarded a housekeeping functional module. Analyzing its components’ expression using two independent data sets (HPA and GTEx), we found for example that *SRPR*, a protein in the signal recognition particle (SRP) receptor, was an extreme gene specific for the stomach, pancreas, liver and salivary gland. This and other examples provide strong evidence on the tuning of a specific component with respect to the average expression of the other components.

While we did not find previous reports on tissue-specific expression and function for most extreme genes, the high load of secreted protein in tissues with many extreme genes is consistent with the idea to provide efficient loading of the secreted proteins on the ER. We also found that PTMs properties specific to a tissue secretome/membrome showed a functional association with the subsystem presenting an extreme gene for that tissue, typically components enriched with PTMs or transport steps. The *PDIA4* gene was an excellent example of this case (Fig. 4B)and (Fig. 7). While *PDIA4*, a specific foldase with disulfide isomerase activity, was a shared extreme gene in the folding subsystem between pancreas, kidney and liver, its expression level was even higher in liver and kidney, consistent with the larger number of secreted proteins with disulfide bonds in these tissues. As we discussed earlier there are many other extreme genes for which their function and existing literatures are strengthening the presence of their tissue-specific fine-tuning, but for the first time we show the evidence for such tuning of specific secretory pathway subsystems. A key question which remains to be explored is whether tissue-specific fine tuning is the result of tissue specialization through evolution or the presence of regulatory programs specific to each tissue to fine-tune the control of its secretory pathway.

**Figure 7.**
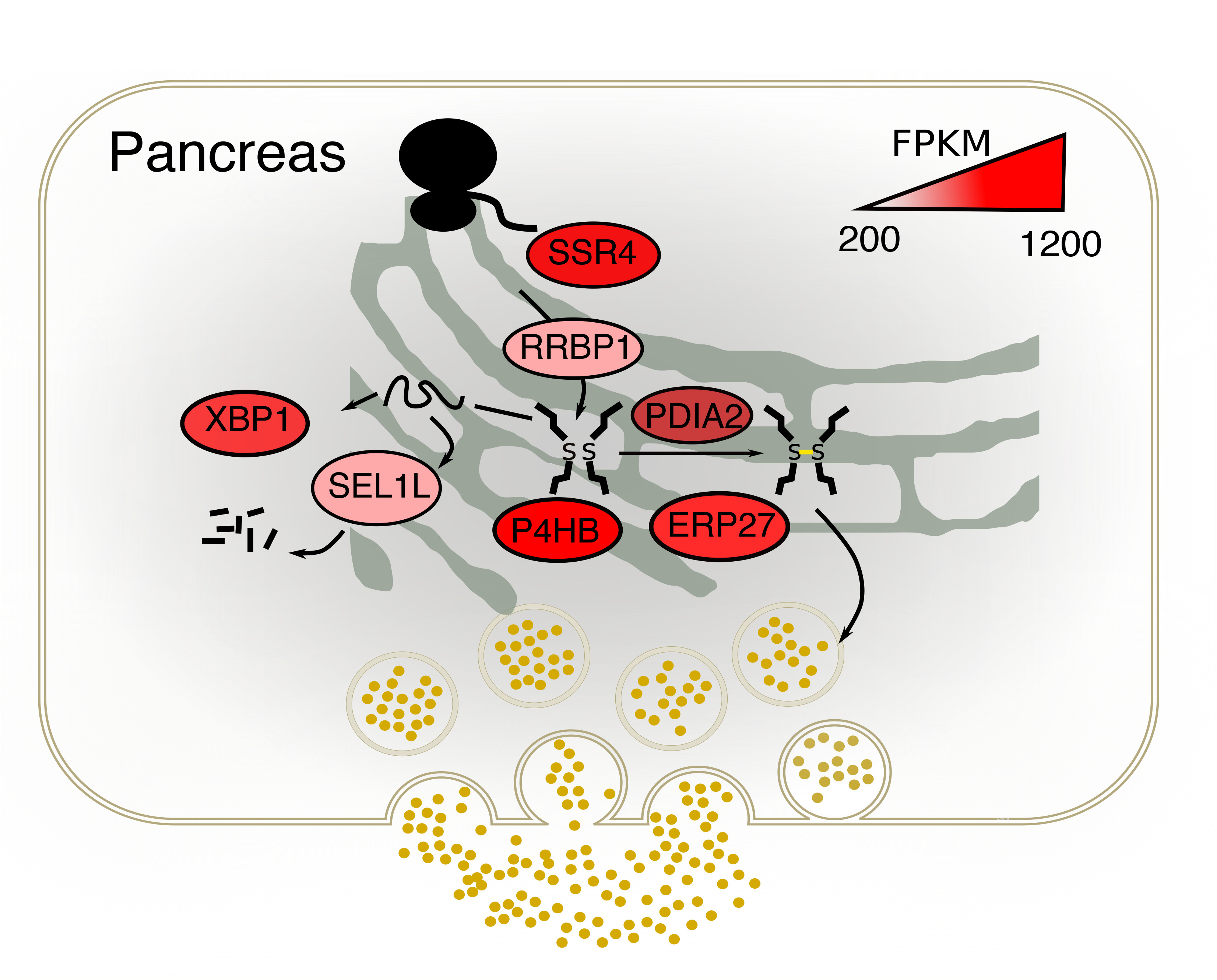
The schematic illustration of pancreas specific extereme genes involved in translocation and folding steps. The detected extreme genes shown as eclipse with their color mapper to their FPKM values in the pancreas. The color for each eclipse is adjusted to the FPKM values (as shown above each object). The genes are involved in translocation subsystem and protein folding (with disulfide isomerase activity) subsystems.

## Supplementary Figures

**Figure S1.**
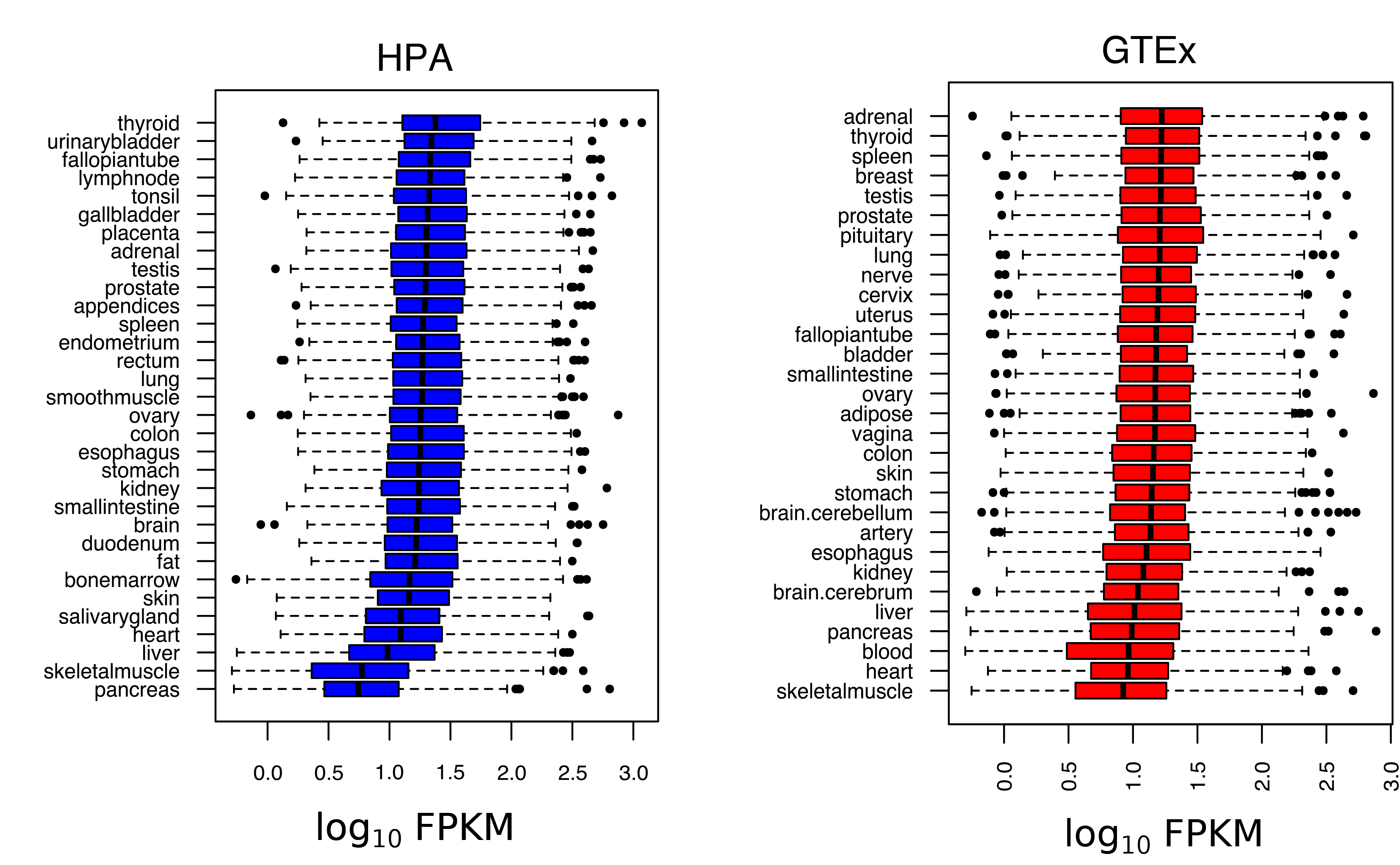
Comparing gene expression distribution of the secretory pathway across human tissues using boxplot in HPA and GTEx data sets.

**Fig S2.**
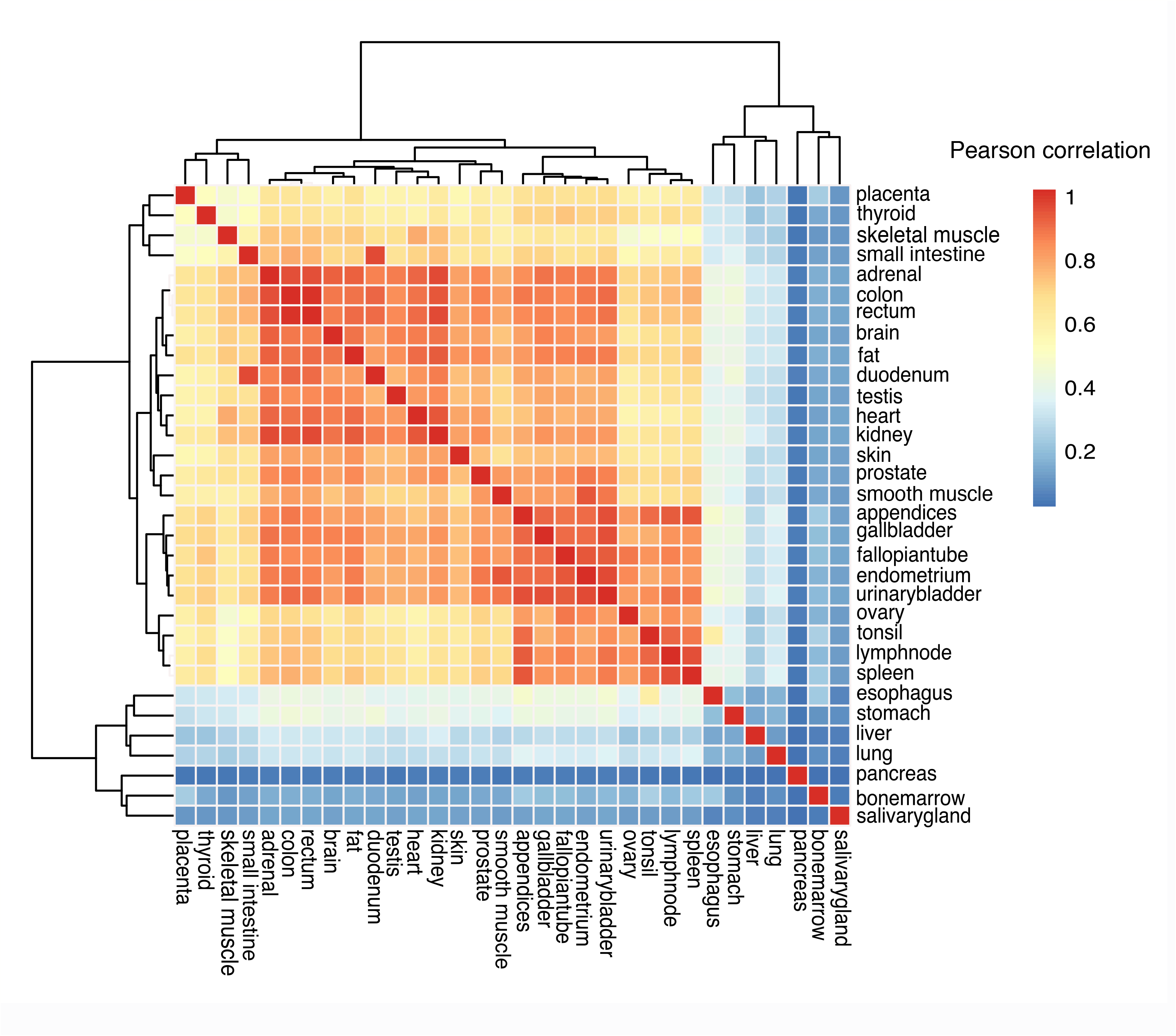
Hierarchical clustering of the secretory pathway components expression profiles obtained for 32 human tissues from HPA data sets. The tissues are clustered based on the Pearson correlation coefficients distance matrix.

**Fig S3.**
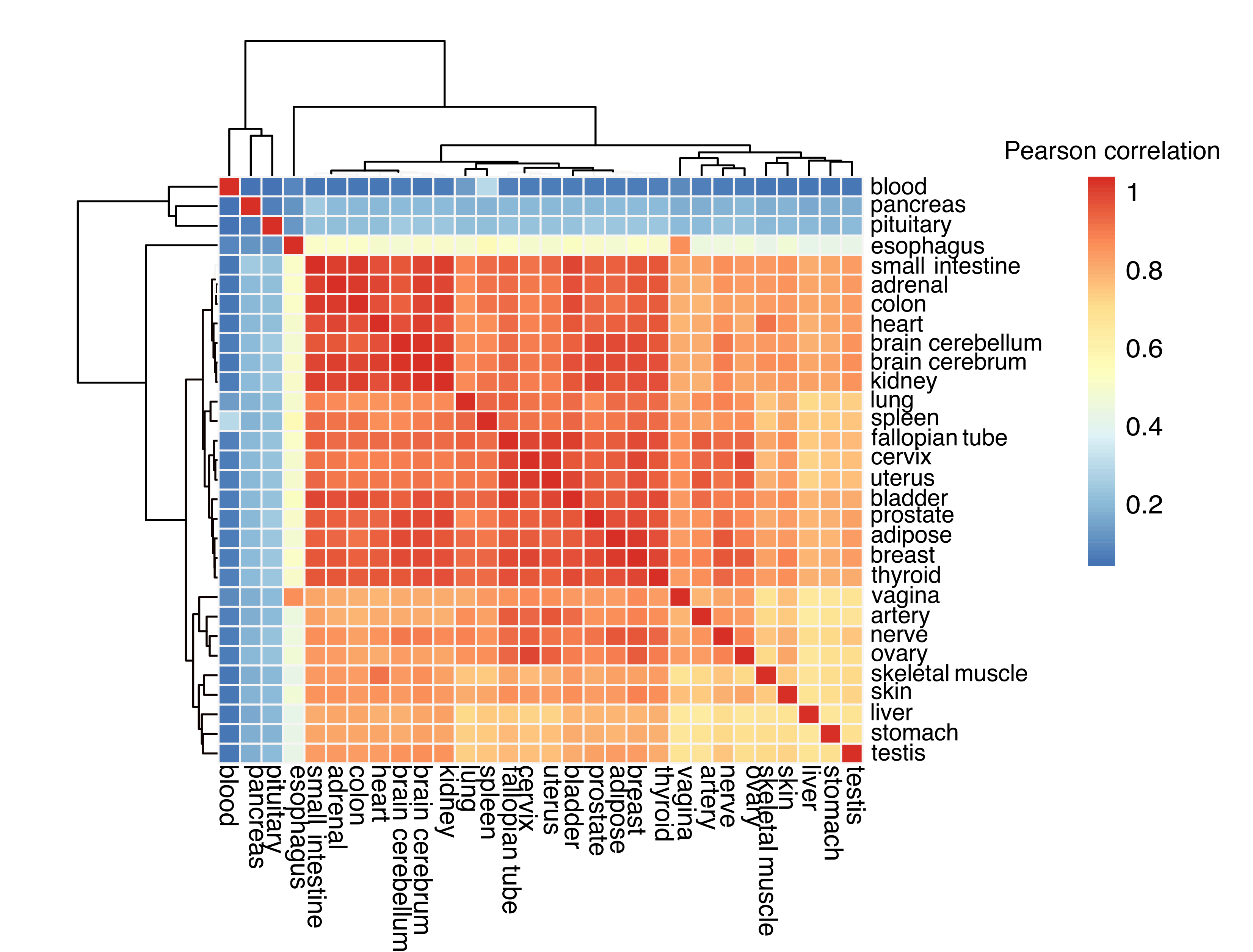
Clustering of the human tissues based on their whole transcriptome data quantified in GTEx data sets.

**Fig S4.**
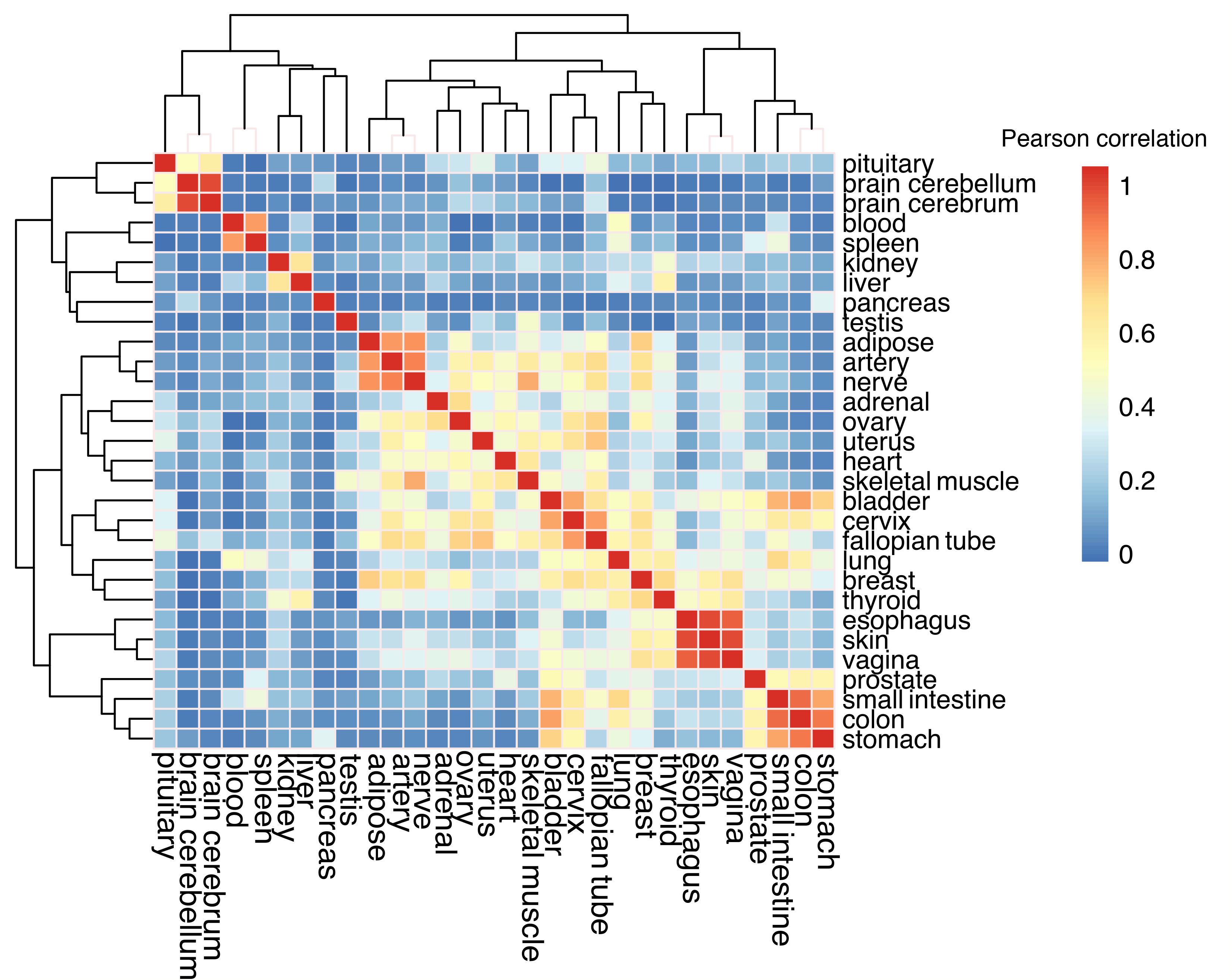
Clustering of the tissues based on genes expression profiles in the tissue-eevated categories obtained from GTEx data sets.

**Fig S5.**
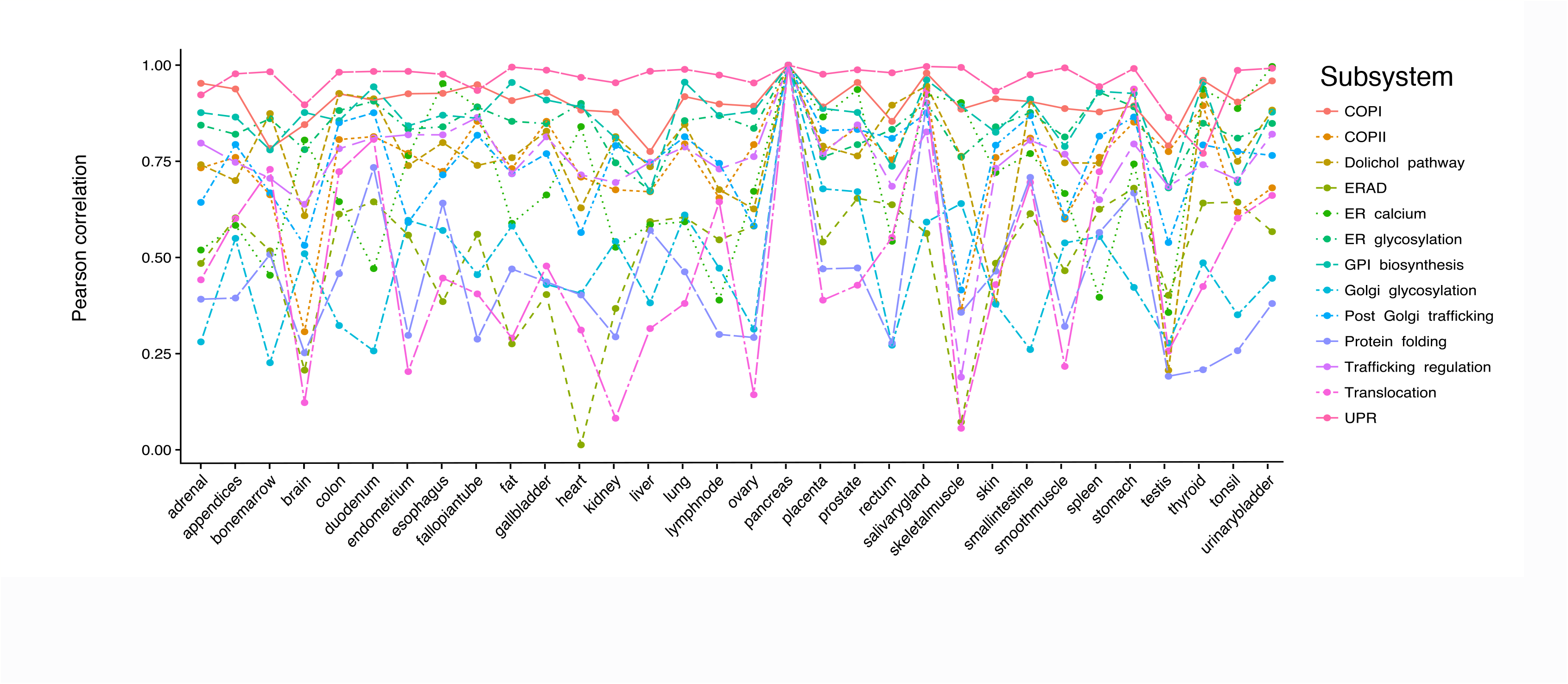
Correlation analysis of pancreas secretory pathway subsystems based on HPA data sets. The multi-line plot displays the correlations coefficients of the gene expression profiles (expressed in all) belonging for each subsystem in pancreas with their counter parts in other tissues. The x- and y-axis in order indicate the tissues and correlations scores. The color code for each subsystem is depicted above the plot.

**Fig S6.**
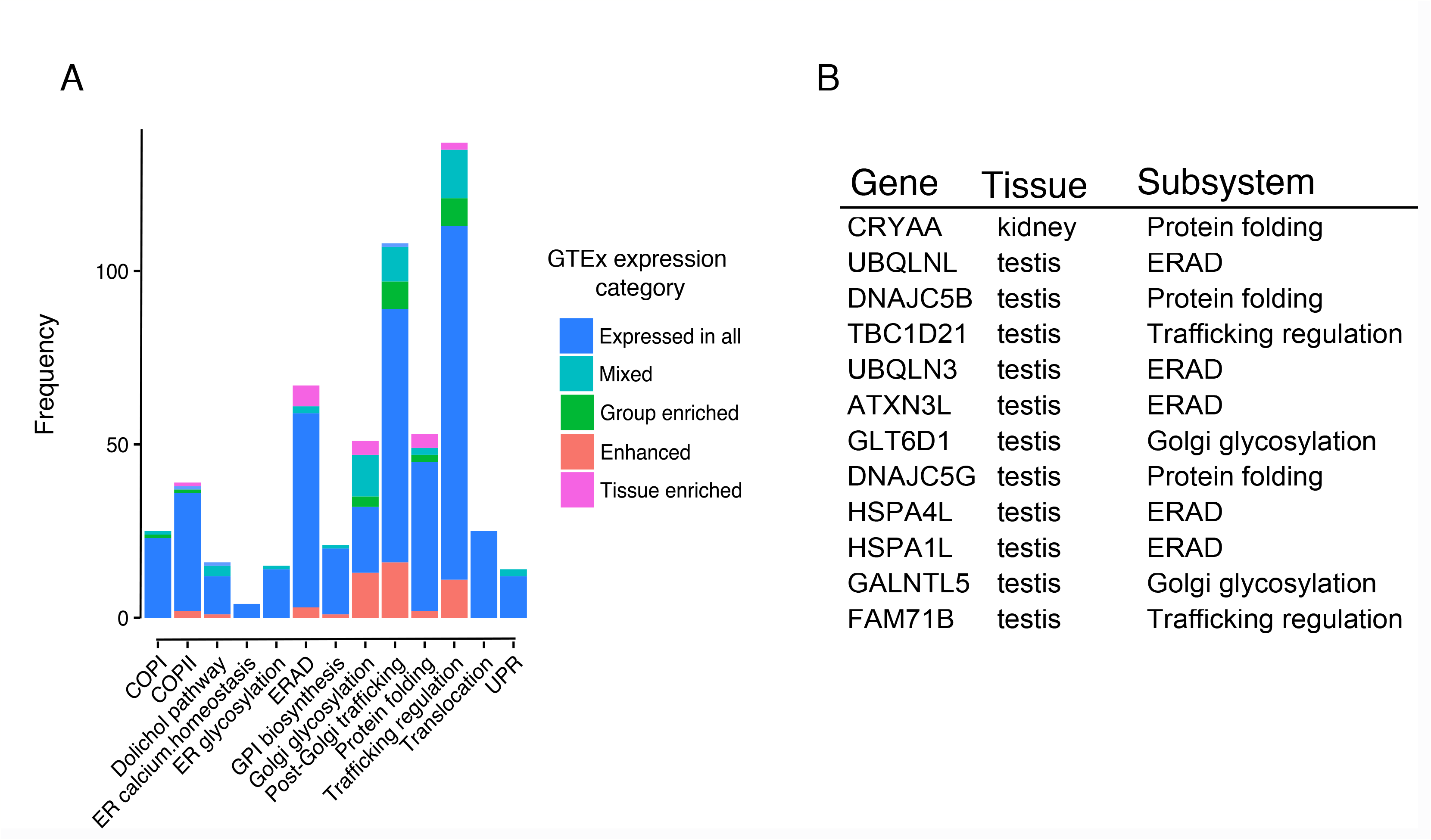
(A) summarizing the secretory pathway genes based on their subsystem and expression categories (B) tissue-enriched genes of the secretory pathway and their corresponding subsystem and tissues.

**Fig S7.**
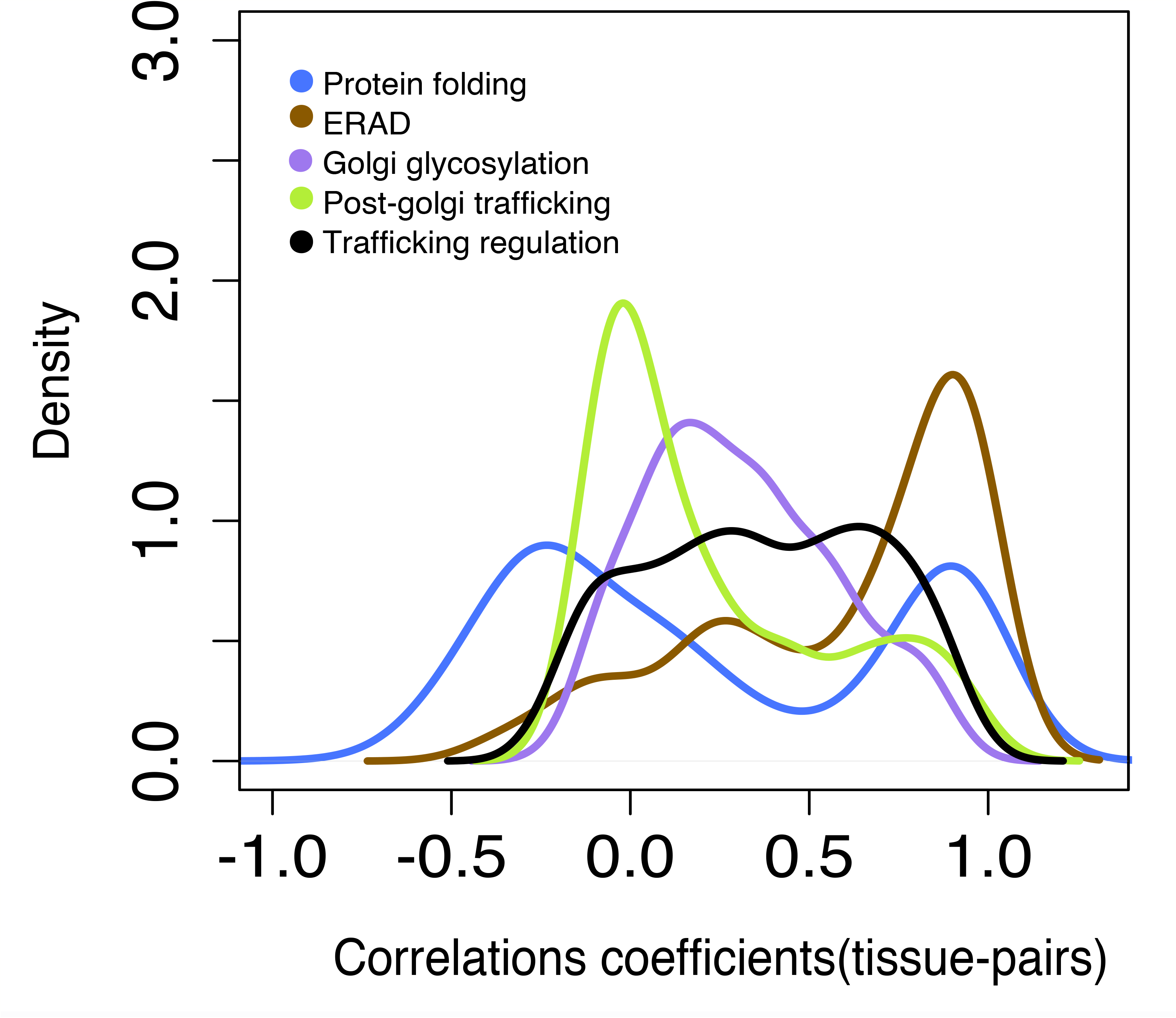
The kernel density plot of the tissue-pairs correlation coefficients (R, x-axis) calculated separately for subsystems having tissue elevated genes (excluding tissue-enriched category).

**Fig S8.**
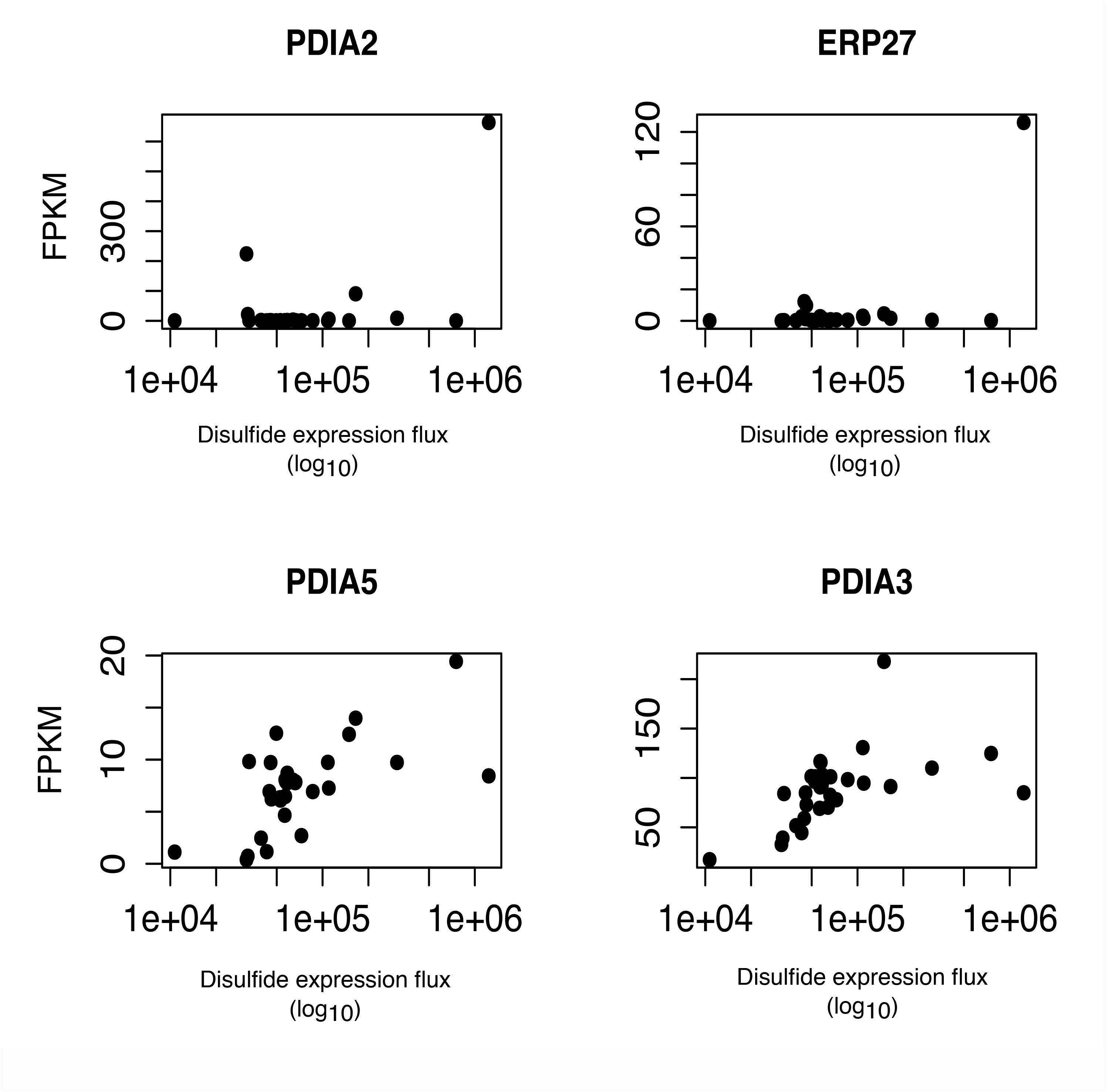
The scatter plots depicting the expression values of the *PDIA2*, ERP27, PDIA5, and PDIA3 in 30 tissues against the total sum of the FPKM values of the tissue-enriched secreted proteins multiplied by the corresponding number of the disulfide bonds of each encoded proteins.

## Materials and Methods

### Transcriptome datasets

We obtained the FPKM values for the human tissues from analysis that has been performed by Uhlén *et al* between (Uhlén et al, 2016) on comparing the recently published RNA-Seq data generated by the Genotype-Tissue Expression (GTEx) consortium(Bahcall, 2015; Melé et al, 2015) and HPA consortium(Uhlén et al, 2015). In these datasets cutoff of 1 FPKM is used to indicate the presence or absence of transcripts for each gene in a tissue. We also used the categorizes defined in their paper. All human protein-coding genes were classified into (i) genes with an elevated expression in one or several tissues, (ii) genes expressed in all analyzed tissues, (iii) genes with mixed expression found in several, but not all tissues, and (iv) genes not detected in any tissues. The elevated genes were further stratified into “tissue enriched”, “group enriched”, or “tissue enhanced”(Uhlén et al, 2016). We used the GTEx data sets as the main expression datasets in our analysis, which its measurements are for 20344 genes across 32 human tissues. The GTEx data is based on measurements for 1641 samples from 175 individuals representing 43 sites: 29 solid organ tissues, 11 brain sub regions, whole blood, and two cell lines: Epstein-Barr virus-transformed lymphocytes (LCL) and cultured fibroblasts from the skin(Melé et al, 2015). The data from HPA(Uhlén et al, 2015) were used in parallel to analyze the consistency.

#### Interactome data

For protein-protein interaction data, we used the CCSB database for humans generated by Rolland *et al* (2014) (Rolland et al, 2014), which includes ~ 14000 high-quality binary protein-protein interactions.

#### Protein Complexes

Protein complex information retrieved from a census of human soluble protein complex data generated by Havugimana *et al* (2012) (Havugimana et al, 2012), which is a network of 13993 high-confidence physical interactions among 3006 stably associated soluble human proteins.

### Data processing, correlation analysis and visualization

we used “plyr”, “tidyr”, and “dplyr” R (https://www.r-project.org/) packages for all data processing steps and correlations analysis. The “pheatmap” and “ggplot2” packages used for visualization of the clustering results and plotting.

### Detection of the extreme genes

To detect the extreme genes in each subsystem we used the Grubbs test(Grubbs, 1950) using “outliers” package in R. The outliers (extreme genes) are collected for all the subsystems across tissues. The Grubbs test assumes the input data has normal distribution, however the gene expression in the subsystems violate this assumption. To avoid the false positives in the detection we run the test based on both HPA and GTEx data and we only select the genes that are detected as outliers using both data sets. The genes also filtered based on the corresponding calculated p-value < 0.05 using standard t-test. The intersect of between the significant extreme genes based on HPA and GTEx are reported as the reliable set of extreme genes for each subsystem for different tissues. The final tissue-extreme genes binary matrix was visualized by pheatmap package in R.

### Secretory pathway network reconstruction

To collect the core components of the human secretory pathway, using the biomart package in R, first we obtained the orthologs of 163 components of our previously reconstructed secretory pathway model in yeast(Feizi et al, 2013). In addition, the additional components were added up to 575, based on comprehensive literature survey and and KEGG secretion-related pathways including protein processing in the endoplasmic reticulum (ko04141) and SNARE interactions in vesicular transport (ko04130)(EV1). By integrating the draft pathway with the CCSB human binary interactome network, we reconstructed a generic network of the human secretory pathway including 15 subsystems (Fig. 1A). The subsystems definition were adopted from our previously work on yeast secretory pathway genome scale model(Feizi et al, 2013).

### Human secretome analysis

#### Defining the human secretome

We parsed the human UniProt GFF file to extract the selected seven secretory features for the human proteome, including the following: *Signal Peptide, N-glycosylation sites, O-glycosylation sites, Disulfide bound, GPI-anchored, Transmembrane domain, Localization*. This information integrated with correlation analysis of the secretome and membrome analysis to find out the tissue specific enrichment of the specific PTMs.

## Acknowledgements

We acknowledge funding from the Knut and Alice Wallenberg Foundation and the Novo Nordisk Foundation.

